# Sex Hormones Alter *Pseudomonas aeruginosa* Iron Acquisition and Virulence Factors

**DOI:** 10.1101/2024.03.12.584604

**Authors:** H Ebrahim, N Bryan, A Collette, S D Armstrong, C Bronowski, R V Floyd, J L Fothergill

**Author notes:** Corresponding author: JL Fothergill.

## Abstract

Urinary Tract Infections (UTI) are one of the most widespread infections in healthcare and community settings worldwide. *Pseudomonas aeruginosa* is the third most common pathogen associated with catheter-associated UTI (CAUTI). *P. aeruginosa* infections are highly resistant and difficult to treat and it is currently classified as priority 1 by the World Health Organisation. *In vitro* studies of microbes typically employ laboratory media. The inadequacy of nutrient-rich media in simulating the physiological environment has led to the development of multiple media that mimic human body fluids, including Artificial Urine Medium (AUM). By studying growth and *in vitro* biofilm assays along with proteomics, we sought to establish whether UTI *P. aeruginosa* respond differently in laboratory media, AUM and urine. To further probe the impact of environmental influences, sex hormones estradiol, progesterone and testosterone were added at physiologically relevant concentrations. The proteomic profiles were then compared between hormone supplemented AUM and urine.

Our findings indicate that bacterial responses in standard laboratory media, AUM and urine were distinct. Increased proteins associated with iron acquisition mechanisms were similar in both AUM and urine. However, differences were observed in other virulence and iron pathways, such as phenazine production. Treatment with hormones decreased the abundance of *P. aeruginosa* proteins involved in iron acquisition. Individual hormones exhibited specific bacterial alterations. The presence of estradiol increased protein abundance of the Pseudomonas Quinolone Signal (PQS) quorum sensing system. This study suggests that *P. aeruginosa* pathogenesis in UTI infections may be influenced by the presence of specific hormones in the host. Understanding the individual role of host factors could contribute to a personalised treatment approach based on the potential impact on infection susceptibility and outcome.

## Introduction

*Pseudomonas aeruginosa* is an opportunistic pathogen that targets immunocompromised hosts suffering from conditions such as cystic fibrosis (CF), wounds/burns, and urinary tract infections (UTIs) (1). The ability of this pathogen to persist in nosocomial settings poses a major challenge to healthcare systems around the world. The World Health organisation has declared *P. aeruginosa* as a priority pathogen that urgently requires new treatments (2). The adaptability of *P. aeruginosa* is attributed in part to its large genome which ranges between 5.5 Mb to 7.0 MB and contains a variety of virulence factors (3).

UTIs are among the most prevalent infections in healthcare and community settings worldwide, affecting approximately 150 million individuals per year (4). UTIs were responsible for 800,000 hospital visits in England (2018-2023), as reported by the National Health Service (5). *Escherichia coli* accounts for approximately 65- 90% of uncomplicated and complicated UTIs (6). Although *P. aeruginosa* infections are not as common, they are often complicated catheter-associated infections (CAUTI) and therefore highly challenging to treat (7). *P. aeruginosa* relies on a combination of biofilm formation and antimicrobial resistance to cause cystitis and pyelonephritis (8). UTIs caused by *P. aeruginosa* are largely understudied, despite their propensity for antibiotic resistance and persistence. *P. aeruginosa* isolates are more likely to be carbapenem-resistant than any other uropathogens (9).

The environmental niches around the body vary and impact bacterial characteristics in different infections. The complexity of the chemical composition within the urinary tract and urine is a challenge, with >4000 different elements and compounds detected in urine derived from both healthy individuals and those with poor health on the metabolome database (10). One variable component in urine are sex hormones. Hormones such as estradiol, testosterone, and progesterone reach peak production in adulthood and diminish gradually as both men and women progress in age (11). There is previous evidence of the impact of these hormones on infectious disease (12–14). A study on the role of sex hormones on clinical outcomes in females with CF found that estradiol induced mucoidy in *P. aeruginosa*. This phenotype is associated with chronic infection and exacerbations were observed in females during the follicular phase, in which estradiol is highest in serum (15). *In vivo* experiments in a murine model found that female mice were more susceptible to respiratory infection by *P. aeruginosa* (13). Sexual dimorphism can also be observed in UTIs; supplementation of estrogen decreased recurrent infections with uropathogenic *E. coli* in post-menopausal women (16,17). Paradoxically, the risk of reproductive-age women developing UTIs is high, partially due to the impact of estrogen (18,19). While the impact of sex hormones on the immune system has been widely reported (20,21), the direct response of *P. aeruginosa* to different sex hormones has not been extensively studied.

In this study, we report the impact of sex hormones estradiol, testosterone, and progesterone on the *P. aeruginosa* proteome in urine mimicking environment. Comparisons between nutrient rich laboratory media, the artificial urine medium (AUM) and pooled human urine were also studied. A focus on the impact of hormones on virulence factors such as iron acquisition mechanisms, quorum sensing and secondary metabolites was performed. Our results show that estradiol in particular, can influence the trajectory of UTI *P. aeruginosa* pathogenesis and therefore may affect outcomes during infection.

## Results

### Clinical UTI *P. aeruginosa* growth characteristics are similar in urine and AUM

In order to study the impact of environment on growth dynamics of *P. aeruginosa*, growth rates of 3 clinical UTI isolates (133043, 133065 and 133098) were measured over a 24-hour period in either AUM, pooled human urine or nutrient rich laboratory media, LB. Isolates grown in LB unsurprisingly showed high growth. In contrast, the isolates in AUM and human urine revealed very limited growth; however, similar levels of growth were observed between the two conditions and was consistently observed across the 3 clinical isolates (Figure 1).

**Figure 1.**
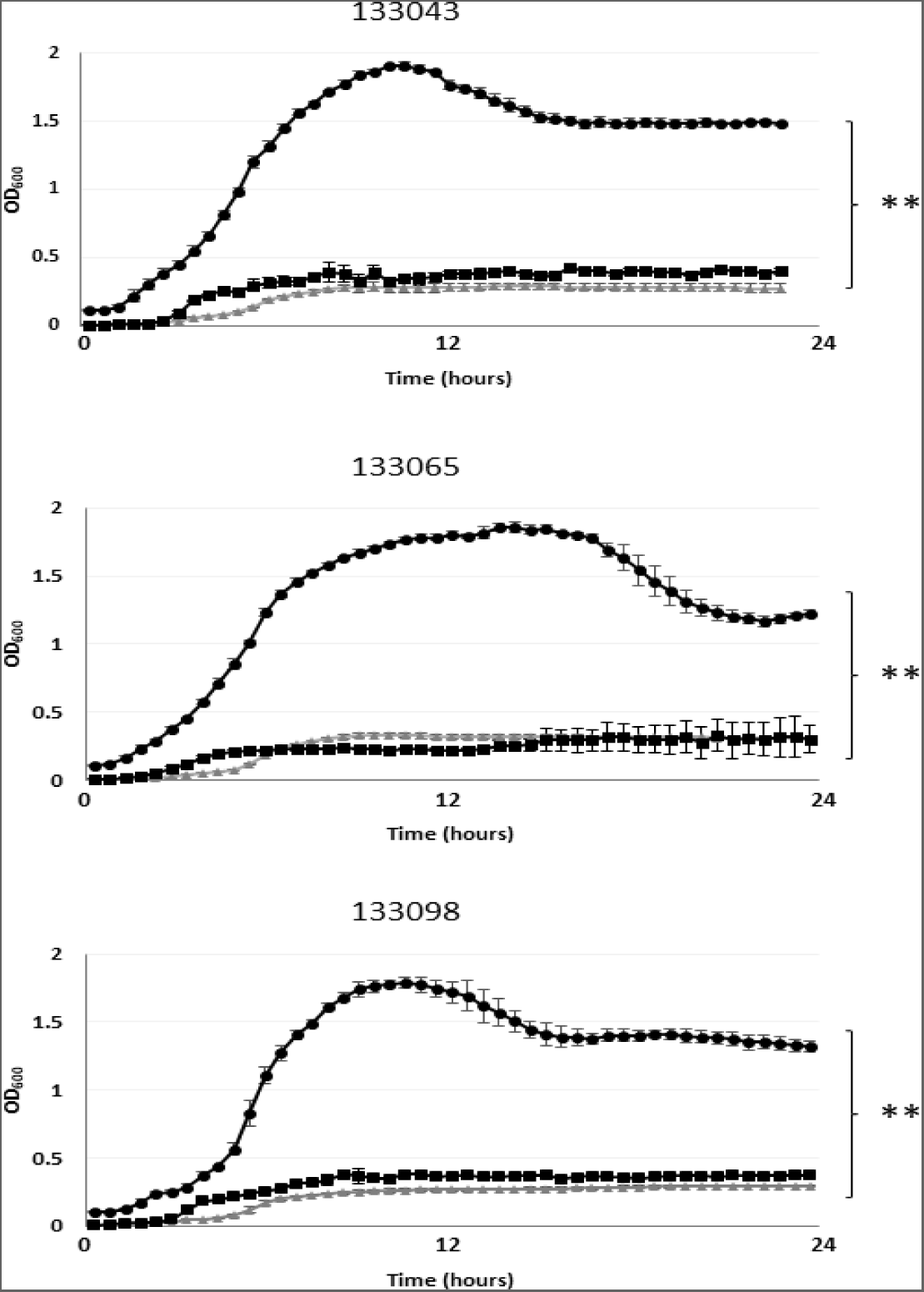
Growth curves of 3 representative *P. aeruginosa* UTI isolates chosen from a panel of 15 UTI isolates in LB broth (black circle) artificial urine media (AUM) (grey triangle) and pooled human urine (black square) over 24 hours at 37°C. Significant differences to the growth in LB are shown using **P<0.001.

Biofilm formation is essential in establishing CAUTI (19). To determine whether there were differences in *P. aeruginosa* biofilm formation when grown in AUM, urine or LB, a panel of UTI isolates and reference strains were subjected to crystal violet staining to determine the biomass under each condition. For all UTI clinical isolates (n=15), the highest attached biomass was observed when grown in LB (Figure 2). All isolates had reduced biofilm in AUM and urine compared to LB at both 24 h and 48 h. To account for differences in growth, the data was analysed to correct for total bacterial growth and expressed as per capita biofilm production (Figure S1). The data shows that under AUM conditions, bacteria had a greater ability to form biofilms compared to LB media, suggesting that proportionally more bacteria (of the total bacteria) adhered to the surface of the plastic in AUM than in LB. AUM conditions may promote biofilm formation.

**Figure 2.**
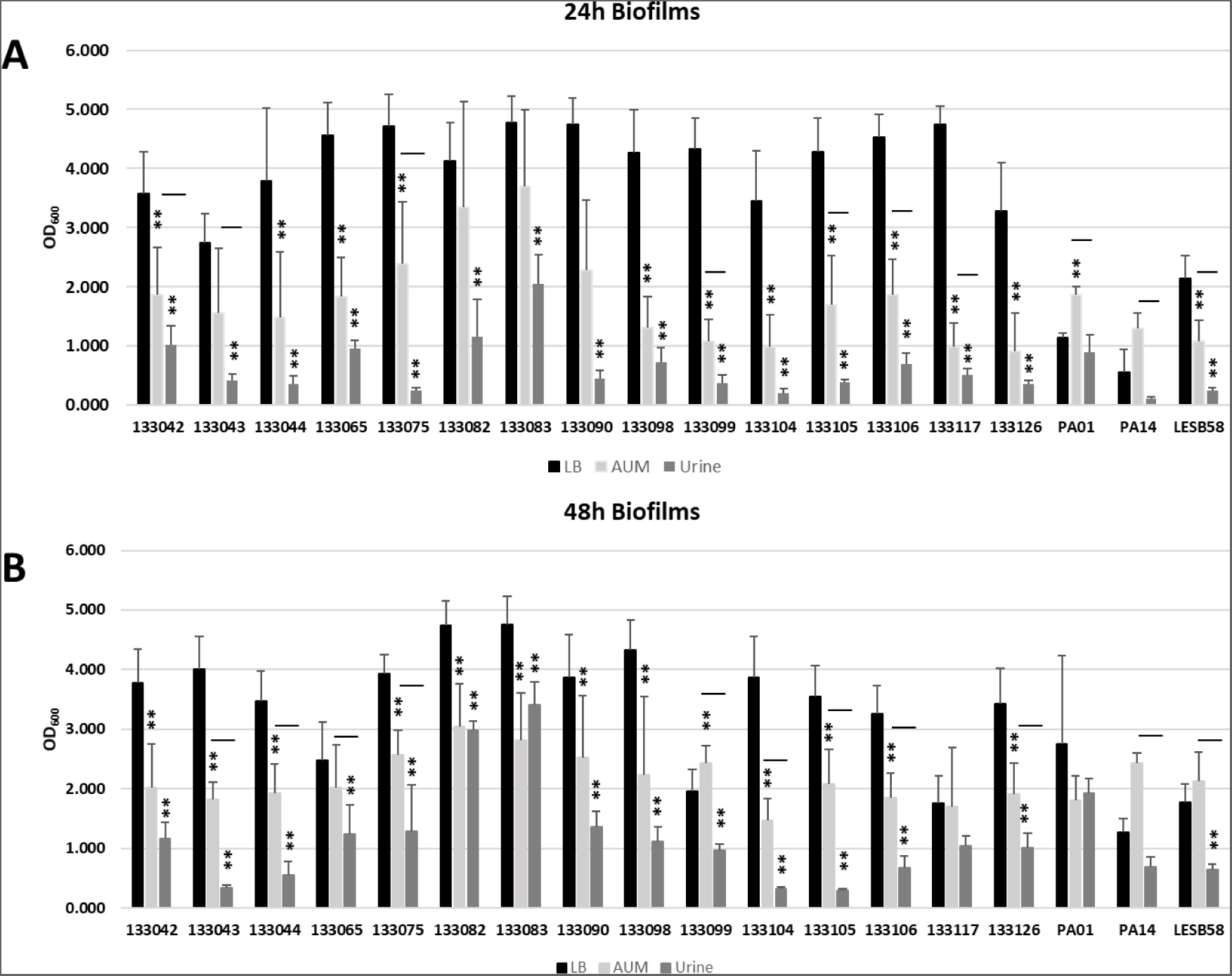
Biofilm assay of *P. aeruginosa* UTI isolates in LB (black), artificial urine media (AUM) (light grey) and pooled human urine (dark grey) after 24 h (Panel A) and 48 h (Panel B). Significant differences to the biomass in LB broth are shown using *P<0.05 **P<0.001; significant differences P<0.05 between AUM and pooled urine are denoted with a solid line.

To investigate this further, a laboratory reference strain (PAO1) and selected clinical isolates were studied using confocal microscopy to study the structure and architecture of the biofilms (Figure S2). Isolates grown in AUM formed denser and tighter biofilms than those grown in the richer nutrient medium, LB. This provides further evidence that AUM promotes enhanced biofilm formation on plastic surfaces.

### Bacteria in AUM and urine display increased abundance of iron acquisition proteins

In order to assess the similarities of *P. aeruginosa* in the artificial urine environment, a single UTI *P. aeruginosa* isolate (133098) was inoculated into LB, AUM, and urine growth media for proteomic analysis.

Variable protein profiles were observed (Figure S3), with 418 proteins displaying increased abundance and 246 proteins showing decreased abundance in both AUM and urine (Figure S4). In AUM, PA2384 showed the most increased abundance compared with the LB control (Figure 3). This protein shows similarity to Fur and has been linked with a large-scale alteration in the expression of proteins associated with iron acquisition and quorum sensing (24). PA2161 was significantly increased in abundance in both AUM and urine. PA2161 is a hypothetical protein that has been previously linked with increased abundance under oxidative stress conditions (25). ExsC showed the greatest reduction in abundance in urine. Given the regulatory role of this protein in the Type III secretion system (T3SS), it seems likely that there is a limited role for T3SS in this environment.

**Figure 3.**
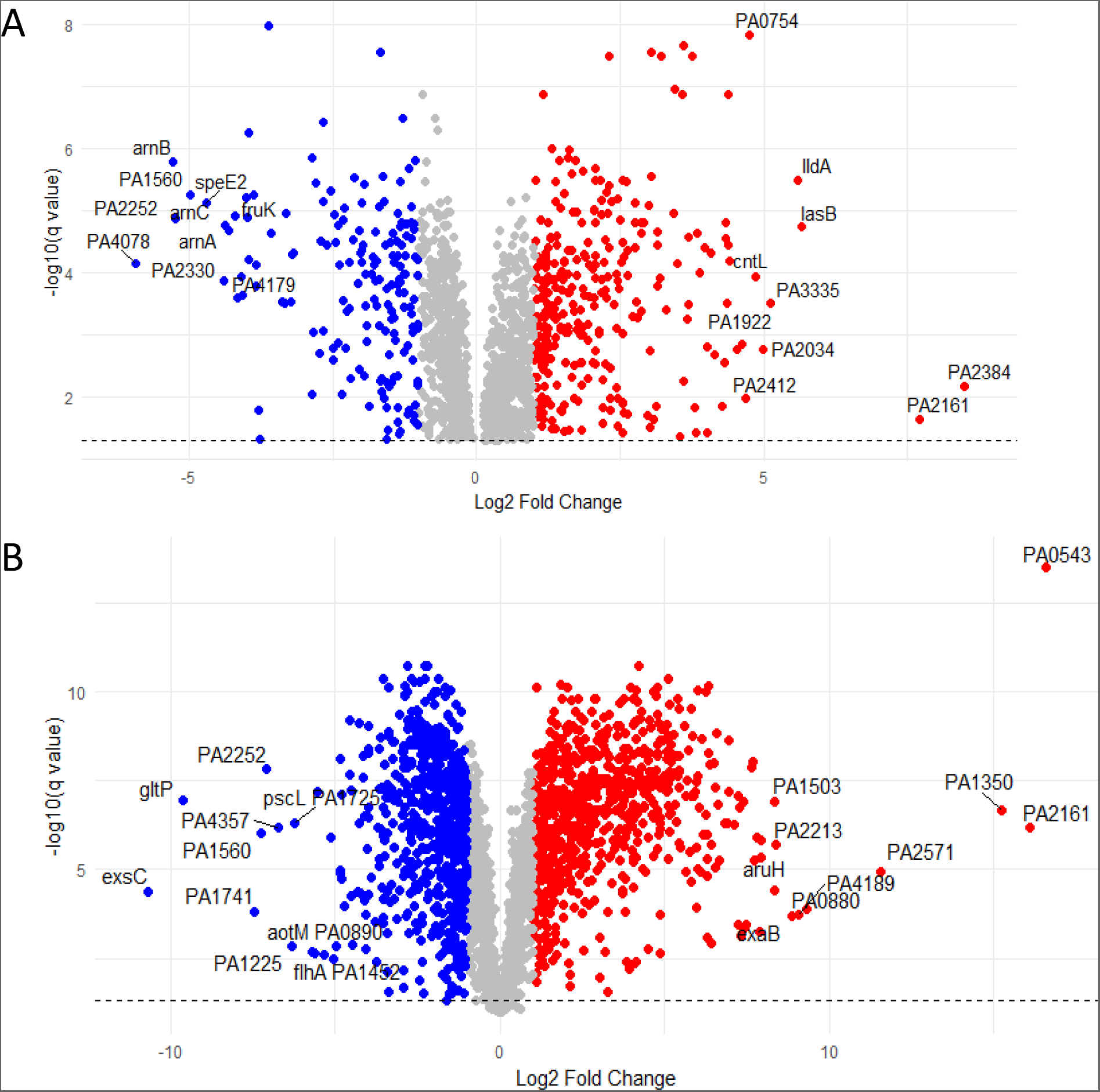
A). Volcano plot of all altered proteins in AUM compared to LB. The top 10 proteins with increased abundance and decreased abundance are labelled. B). Volcano plot of all altered proteins in urine compared to LB. The top 10 proteins with increased abundance and decreased abundance are labelled. Significance refers to the q value determined by five replicates in each condition. Proteins with a significant q value (<0.05) are shown in red for those with increased abundance and blue for those with decreased abundance.

In both AUM and urine, higher protein abundance was observed in iron acquisition proteins, particularly in the pyoverdine (Pvd) pathway (Figure 4). With the exception of PvdE, 11/12 Pvd proteins detected displayed increased abundance in AUM and urine compared to LB. Notably, PvdN, PvdO, and PvdP were produced at 31, 35, and 16-fold increased levels in urine than in LB, respectively. FpvA, a TonB-dependent receptor (TBDR) that transports ferric- pyoverdine substances into the periplasm of the bacterial cell (26) was also found in increased abundance in both AUM and urine. In this instance, FpvA growth in urine resulted in a 15.4- fold increase and a 3.7-fold increase in AUM (Figure 3). This indicates that, in terms of changes in pyoverdine-related proteins, *P. aeruginosa* grown in AUM and urine display similar responses. The abundance of proteins involved in pyochelin production displayed a more variable response. However, >1000 proteins displayed altered abundance in urine that were not captured within AUM. This highlights the complexity of urine.

**Figure 4.**
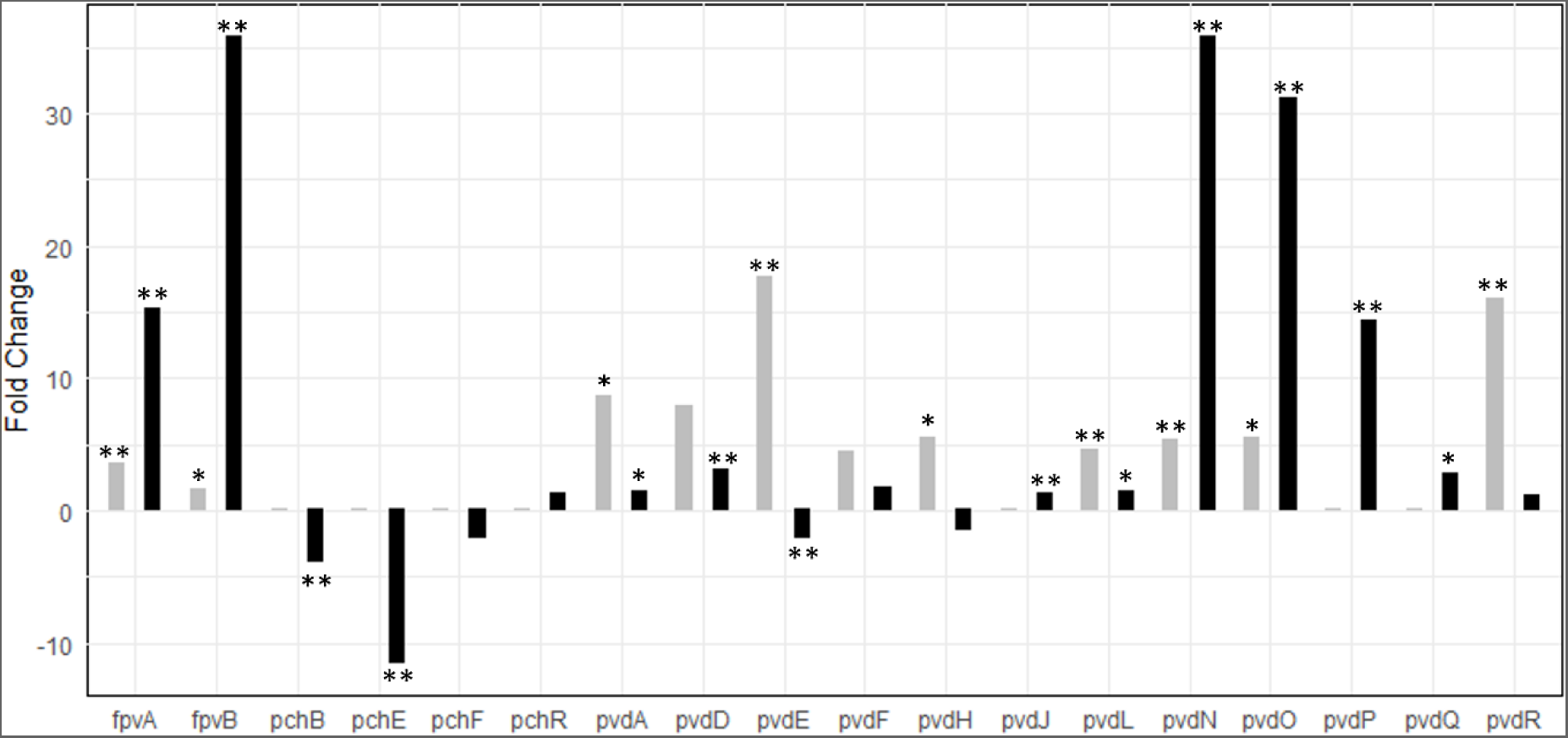
Production of iron acquisition proteins in abundance in AUM and urine relative to LB. * denotes a q-value ≤ 0.05; ** denotes a q-value ≤ 0.01.

### Sex hormones alter the *P. aeruginosa* proteome

In order to assess the abundance of proteins following *P. aeruginosa* growth with estradiol, testosterone, and progesterone in AUM, a single clinical isolate (133098) was chosen for further study. For estradiol, 127 proteins were significantly increased and 93 were significantly decreased in abundance compared to the vehicle control (AUM-V) (Figure S5). For progesterone, 180 proteins were significantly increased and 155 were significantly decreased in abundance, and for testosterone, 138 proteins were significantly increased and 165 were significantly decreased in abundance compared to the control (Figure 5). The proteins with the highest abundance compared to the control were PA4131, AmiE, and PA0122 for estradiol, progesterone, and testosterone, respectively.PA4131 is a probable iron-sulphur protein, and AmiE is an aliphatic amidase, both influenced by hydrogen cyanide production(27,28). PA0122 (RahU) plays a role in innate immunity modulation (29). The proteins with the lowest abundance were PA5487, PchG, and PA2550 for estradiol, progesterone and testosterone, respectively. PA5847 (DghC) is a diguanylate cyclase, involved in cyclic di-GMP production (30). PchG is involved in the synthesis of pyochelin, and PA2550 is a probable acyl Co-A dehydrogenase (31).

**Figure 5.**
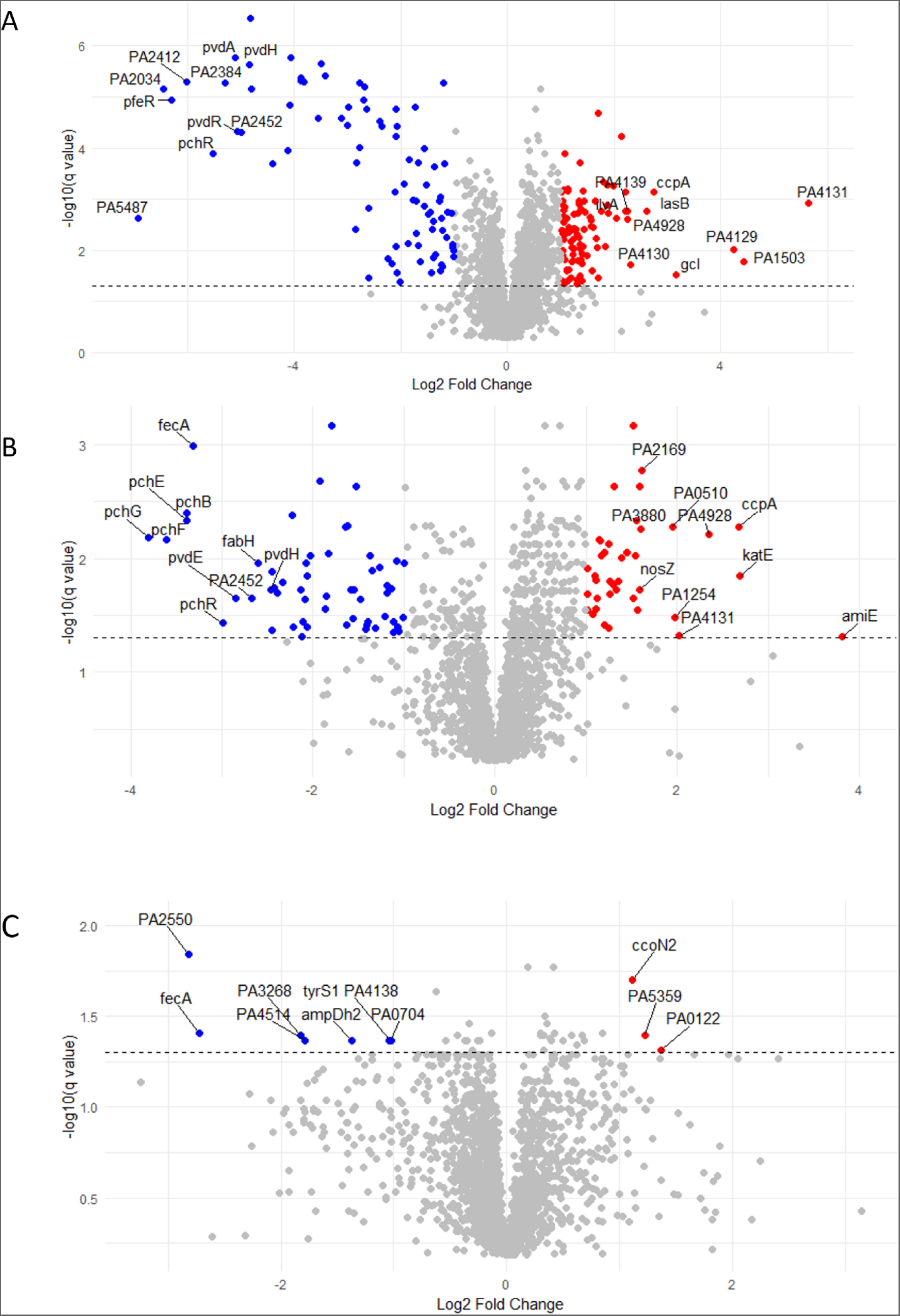
Altered *P. aeruginosa* protein abundance in the presence of sex hormones. A) Volcano plot of all altered proteins in the treated samples with estradiol compared to AUM-V. B) Volcano plot of all altered proteins in the treated samples with progesterone compared to AUM-V. C) Volcano plot of all altered proteins in the treated samples with testosterone compared to AUM-V. The blue dots represent down-regulated differentially expressed proteins, the red dots represent up-regulated differentially expressed proteins, and the grey dots represent proteins that are differentially expressed; however, the differences are not significant.

### The impact of sex hormones on iron acquisition mechanisms

Iron acquisition mechanisms are indispensable in *P. aeruginosa* infections (26). The impact of sex hormones on these mechanisms was analysed. Pyochelin is a siderophore produced to counteract iron scarcity in the host. All proteins involved in pyochelin biosynthesis were less abundant across all hormone conditions compared to AUM (AUM-V, the vehicle control), and these were decreased in abundance from −2 to −8-fold compared to the control. This impact on siderophore-related proteins was also identified for those in the pyoverdine synthesis pathway, where synthase enzymes and receptor proteins were less abundant in the presence of hormones. Heme uptake and other proteins associated with iron acquisition were also less abundant than AUM-V (Figure 6).

**Figure 6.**
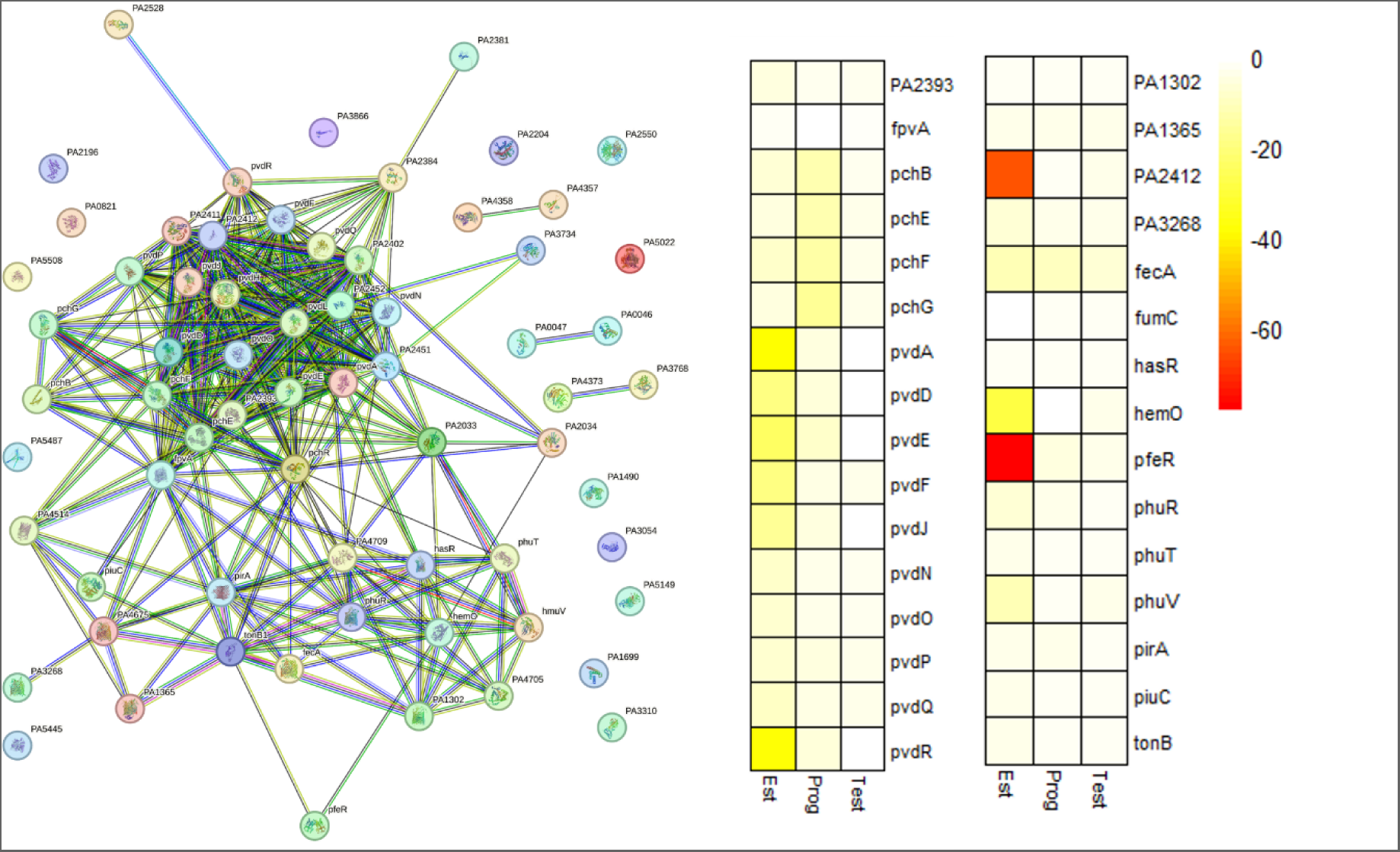
Impact of sex hormones on iron acquisition proteins. A). String plot of all proteins with decreased abundance in the presence on estradiol showing a major cluster of proteins involved in iron acquisition (KEGG enrichment - Biosynthesis of siderophore group nonribosomal peptides p=0.00068). B) Summary heatmap of 31 iron acquisition proteins that were detected with altered abundance in the presence of estradiol, testosterone, and progesterone.

*P. aeruginosa* can also hijack siderophores made by other bacteria. ChtA and FecA contribute to the capture of xenosiderophores and were found to be less abundant in all hormone treated AUM (Figure 6) (26). Furthermore, PirA is a receptor for ferrioenterobactin and shows a 9 -fold decrease in abundance. Expression of this receptor is under the control of the PfeRS two-component system (25), and the PfeR response regulator was downregulated by 78-fold in the estradiol samples and to a lesser extent in the presence of progesterone and testosterone (5-fold and 3-fold reduction, respectively).

Other proteins were also decreased, such as fumarate hydratase FumC, which is produced in response to iron starvation and also linked to pyochelin and pyoverdine production (32). PA2033, a hypothetical protein that has been linked to a novel iron-acquisition mechanism (33), was found to be decreased by estradiol, testosterone, and progesterone by approximately 6-fold, 2-fold, and 4-fold, respectively. There is a clear impact caused by all three hormones, leading to a decreased abundance of proteins associated with iron acquisition. Taken together, the presence of each of the three hormones results in a globally decreased abundance of proteins associated with iron acquisition. This impact was strongest in the presence of estradiol.

### Estradiol is associated with increased abundance of the Pqs system and secondary metabolites

Production of PQS-associated proteins was increased in estradiol compared with progesterone and testosterone-supplemented AUM (Figure 7). A >2-fold increase in comparison to AUM-V and PqsA-E was observed in estradiol. The PQS system has been implicated in the control of the production of phenazines. Along with increases in PQS proteins, an increase in 9 proteins linked to phenazine biosynthesis was observed. Estradiol appears to have a direct impact on *P. aeruginosa*, leading to an increased abundance of proteins associated with the PQS system, including phenazine biosynthesis. This effect was not observed in the presence of the other two hormones, suggesting differential effects based on hormone identity.

**Figure 7.**
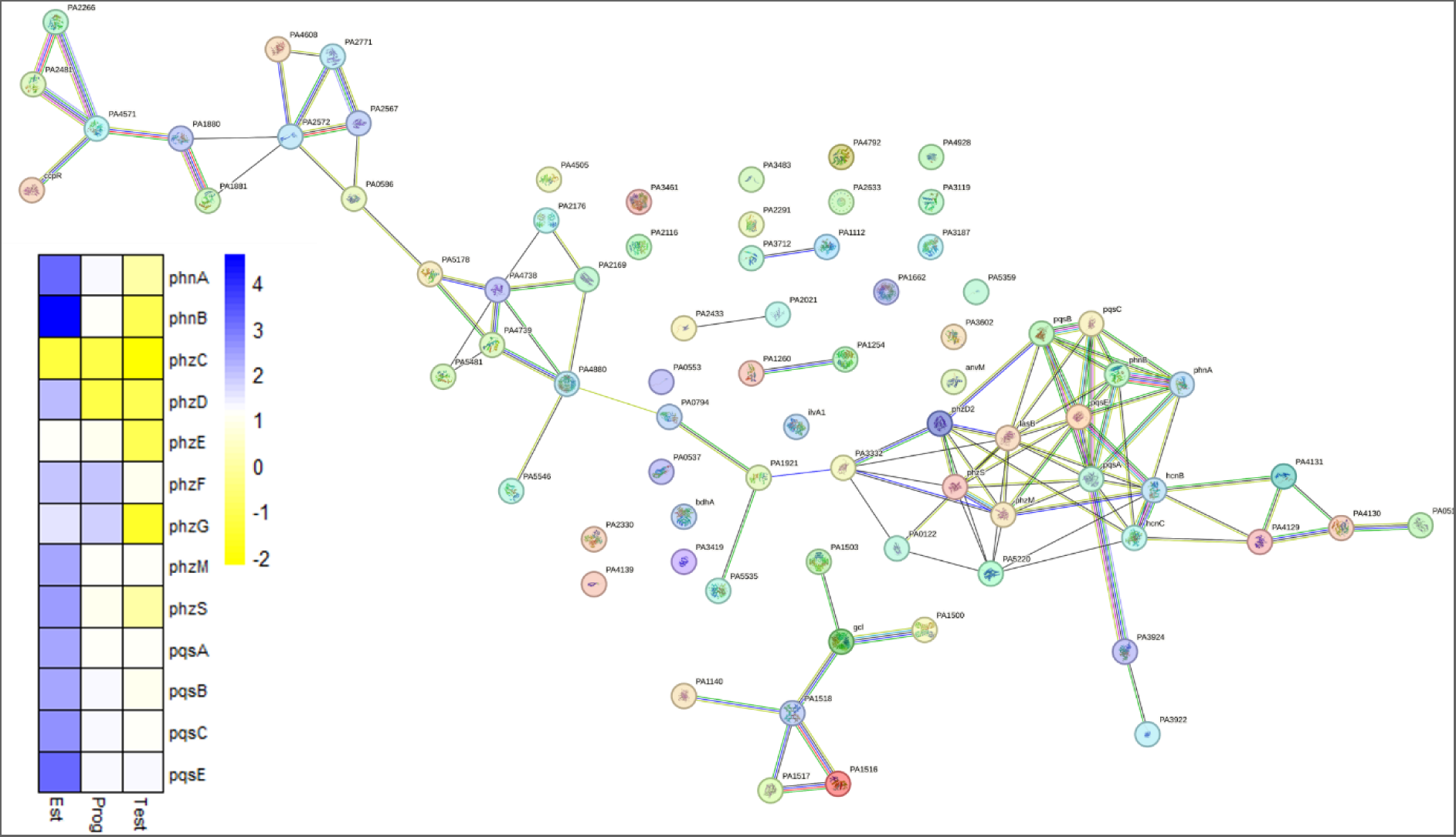
Abundance of *P. aeruginosa* proteins associated with the PQS system, phenazine biosynthesis. A). String plot of all proteins with increased abundance in the presence on estradiol showing a major cluster of proteins involved in phenazine production (KEGG enrichment Phenazine biosynthesis – p=9.73e-09, Quorum sensing p=0.0029 and biofilm formation - 0.0377). B). Summary heatmap of 13 phenazine and Pqs quorum sensing proteins that were detected with altered abundance in the presence of estradiol, testosterone, and progesterone. Increased abundance is shown in blue and decreased abundance is in yellow.

The presence of estradiol was also associated with increased levels of a key virulence factor, LasB, with a 6.1-fold increase compared to the control (Figure 8). LasB has been implicated in niche establishment, overcoming host immune responses and biofilm formation that could be beneficial in persistent infections (34).

**Figure 8.**
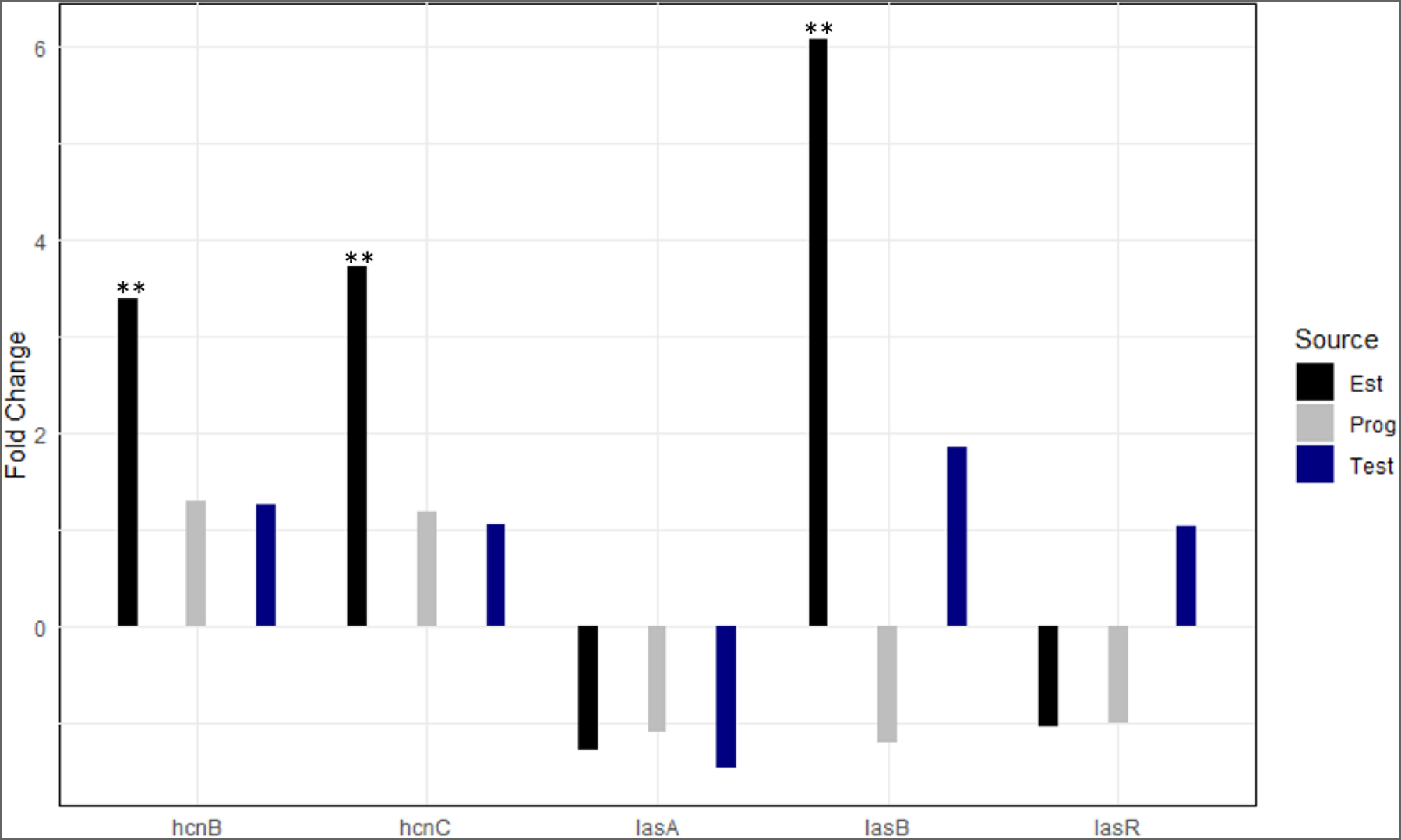
Protein fold change of proteins involved in the las quorum sensing system and virulence factors LasB (elastase) and proteins associated with hydrogen cyanide production (HcnB and HcnC) in *P. aeruginosa* grown in the presence of hormones. Fold change of the protein abundance produced by *P. aeruginosa* in AUM containing either estradiol, testosterone, or progesterone compared to an AUM-vehicle control is shown. A significantly increased abundance (>1.5-fold change and significant q-value) is only observed in the presence of estradiol (eg. LasB displays a 6-fold change in comparison to AUM-containing vehicle control). ** denotes a p-value ≤ 0.01.

## Discussion

Sex hormones play a crucial role in modulating many physiological processes in humans. While studies have previously linked hormones to altered immune responses, less focus has been given to the direct impact of hormones on bacterial infections. The aim of this study was to establish the influence of specific sex hormones on *P. aeruginosa in vitro* in conditions mimicking the environment of the urinary tract.

Initially, we established the similarities in the proteome between AUM, urine, and standard laboratory media. Each medium clustered separately using principal component analysis, and a large number of proteins were differentially altered in urine. Urine is a highly complex biological medium (35). The proteins with the highest abundance compared to LB were hypothetical proteins, PA2161 and PA1350. Little is known about their potential role, particularly in UTIs. ExsC, part of the T3SS regulatory system, displayed the greatest decrease. A decrease in this anti-activator is likely to result in decreased activity of the T3SS. The T3SS has been implicated in the acute phase of UTIs but decreased in the chronic phase (36).

AUM is a simplified *in vitr*o model that mimics the urinary environment. While our data suggest that many factors that can influence bacterial characteristics in urine may be missing in AUM, some key pathways were similar in both environments. One of the iron acquisition systems is the pyoverdine system, which is synthesized in response to severe iron starvation (26). When *P. aeruginosa* encounters environments with iron scarcity, FpvA initiates a signalling cascade that turns on the pvd and pyochelin (pch) genes. (37). While pyoverdine pathway proteins showed similar increases in abundance compared to LB, some proteins (FpvB, PvdN and PvdO) were produced in higher quantities in urine than AUM. While AUM may lack the complexity of urine, we show here that there are similarities in key pathways associated with infection and virulence.

AUM was used as a base to study the direct impact of different sex hormones (estradiol, testosterone, and progesterone). While some responses were hormone-specific, key similarities could be seen in the bacterial response to all three hormones. The pyochelin pathway displayed decreased abundance in the presence of estradiol, progesterone, and testosterone. *P. aeruginosa* utilises pyochelin when iron is less limited in the surrounding environment relative to the state in which pyoverdine is produced and utilised (38).

Furthermore, our results show that proteins involved in haem extraction from the host are significantly reduced. Haem uptake from the host is conducted via two different pathways, the Phu and HasR systems (39). Further reductions of intracellular proteins involved in processing haem, such as PhuS and the heme oxygenase, were observed. All sex hormones in this study reduced fumarase C, a protein that is part of the tricarboxylic acid cycle. This protein is produced in response to iron starvation and is involved in alginate biosynthesis and conversion to mucoidy (32).

*P. aeruginosa* can also hijack siderophores made by other microorganisms (26). All sex hormones in AUM reduced the expression of proteins ChtA, PfeR and PfeS. This could potentially impact *P. aeruginosa* ability to obtain xenosiderophores in conditions mimicking CAUTIs. This reduction may be due to greater availability of iron, thereby indicating that in the presence of hormones, iron limitation is less severe and the bacteria may experience reduced nutritional immunity (40). However, the mechanism underlying this process is unknown. Alternatively, hormones could directly reduce the production of these iron acquisition proteins. As a result, other pathogens may have an advantage over *P. aeruginosa* during polymicrobial infections in accessing iron. In a study which utilised proteomics to assess the response strain PAO1 to iron starvation, all classical iron upregulation proteins of haem, pyochelin and pyoverdine were upregulated (41). This is in contrast to our results, which show that iron acquisition mechanisms are downregulated. Therefore, this may support the suggestion that the bioavailability of iron in AUM containing hormones is increased.

Quorum sensing (QS) mechanisms control the production of virulence factors such as phenazines and elastases. PQS is the third QS system identified in *P. aeruginosa* and can promote the synthesis of phenazines (42). In this study, PQS proteins were higher in estradiol-treated samples. PQS is associated with higher virulence, as shown in an acute UTI mouse model (43). *P. aeruginosa* with a fully functional PQS biosynthetic pathway resulted in higher virulence, tissue destruction, and severe inflammatory responses in mice (43). Estradiol increased the abundance of all phenazine biosynthesis proteins. Phenazines, in particular pyocyanin, may have a variety of effects during UTIs. Pyocyanin may impair host cell repair and induce inflammation. It has also been implicated in biofilm formation and the production of damaging reactive oxygen species (7).

In addition to pyocyanin, another virulence factor (LasB) also showed higher abundance in the presence of estradiol. LasB has been reported to play a role in immunomodulation, biofilm formation, and elastylatic activity (34). Attenuation of LasB is an attractive target for therapeutic as targeting a virulence factor would likely circumvent the selective pressure that can drive the emergence of antibiotic resistance. Hydrogen cyanide (HCN) biosynthesis protein production was also increased in the presence of estradiol. Furthermore, PA4131 showed the highest increase in estradiol and this has been shown to be induced by HCN and linked to enhanced survival in the presence of this toxin (28). These data highlight that estradiol, in particular, may have a combinatorial impact on virulence.

Estradiol has been linked with exacerbations of lung infection in women with CF. This could be a contributing factor to the worsened disease outcomes in women compared to their male counterparts and earlier mortality (15,44). This study provides further insight into the direct impact of estradiol on *P. aeruginosa* infection; however, further work is needed on the clinical and longitudinal impact of sex hormones on UTIs. The use of oral contraceptives containing estrogen-progestin to prevent pregnancy in premenopausal women affects these levels and therefore could be explored as an intervention to alter bacterial pathogenesis during infections, particularly for those at increased risk or with persistent infection (15,45).

This study demonstrates that sex hormones can have a consistent effect on iron acquisition systems. However, other systems may be affected exclusively by a single hormone. These modifications may have implications for the pathogenesis of *P. aeruginosa* in UTIs. The production of reproductive hormones decreases with age in both male and female patients. Thus, a greater understanding of hormone-dependent host-pathogen interactions is necessary and may contribute to the development of a personalised medical approach.

## Methods

### Isolate storage and culture

A single *P. aeruginosa* colony was used to inoculate LB broth (Sigma Aldrich), artificial urine medium (AUM) or pooled human urine in glass or plastic universal tubes overnight. All cultures were grown at 37°C and shaking at 180 rpm unless otherwise stated. LB agar (Sigma Aldrich) was used to grow bacterial cultures, which were spread inoculated onto the agar and allowed to grow overnight at 37 °C.

### Artificial urine medium (AUM preparation)

AUM was prepared using distilled water with the basic components as formulated by Brooks *et al,* (1997) (46).

### Collection of pooled urine

Pooled urine was collected as per ethical guidelines of the University of Liverpool from 2 healthy males and 2 healthy females with consent. This was conducted by mixing equal amounts of urine after capturing midstream urine. The mixed urine was then filtered with a 0.2 µM vacuum filter into a sterile Duran bottle. Maximum time period for use was 10 days.

### Biofilm assay (Crystal Violet staining)

The 15 clinical isolates’ overnight cultures were diluted 1:100 in LB or AUM. Then, 200 µl of each *P. aeruginosa* -containing LB or AUM solution was added in quadruplicate to a 96-well Corning® Costar® plate and grown at 37°C for 24h or 48h. The wells were subsequently washed by PBS and washed with crystal violet (CV. The wells were washed and solubilized with 95% of ethanol (Sigma Aldrich) prior to the biomass measurement at OD600nm by the Floustar Omega plate reader. Laboratory reference strains PAO1 (47) and LESB58 (48) were included as controls.

### Biofilm microscopy

For biofilm microscopy, biofilms were prepared using glass microscopy-compatible plates (Greiner high and medium binding 96 well plates, Sigma-Aldrich) according to the method described above. Biofilms were allowed to grow at 37°C for 24 hours. The media and planktonic *P. aeruginosa* cells were removed without disrupting the well’s bottom. Each well was washed gently with LB and then removed. 3.58 ml of PBS was combined with 5 µl of two dyes from the LIVE/DEAD® BacLight® Bacterial Viability Kit (Thermo Fischer Scientific): SYTO 9 dye (3.34 mM) and Propidium iodide (20 mM). Only viable cells/biofilms were considered for analysis. 50 µl of the dye solution was used to stain each well. After 30 minutes, biofilm microscopy was conducted. To image biofilms in Z-stacks, a Carl Zeiss confical microscope with a 40x oil lens was utilised.

### Growth curves

To determine the rate of bacterial growth. Overnight overnight cultures were diluted at a ratio of 1:100 in polystyrene 96-well plates manufactured by Corning® Costar®. The experiment was conducted using five technical replicates and five biological replicates. Growth curves were performed in order to determine the optical density in each media and continuously assess growth. Once the optical density of the cultures reached 0.25 +/- 0.05, the proteome of the candidate bacterial strain was isolated.

### Protein extraction

Briefly, the bacterial cell pellets from candidate isolate 133098 were lysed by the addition of 1% (w/v) sodium deoxycholate in 50 mM ammonium bicarbonate, followed by sonication. The total protein content of the clarified lysate was measured using the Pierce™ Bradford Protein Assay Kit . Protein concentrations were normalised before reduction and alkylation of cysteines. The samples were then incubated with proteomic grade trypsin (Sigma) overnight before acidification with TFA to a final concentration of 0.5% (v/v). The digests were centrifuged at 12,000 x g for 30 min to remove precipitated sodium deoxycholate.

### NanoLC ESI-MS/MS peptide analysis

Peptides were introduced to the mass spectrometer using a Ultimate 3000 HPLC system (Dionex/Thermo Fisher Scientific)equipped with an Easy-Spray PepMap® RSLC column(50 cm × 75 μm inner diameter, C18, 2 μm, 100 Å). The column was kept at a constant 35°C. Peptide separation was performed with 0.1% formic acid (Buffer A) and 80% acetonitrile in 0.1% formic acid (buffer B), using a linear gradient of 3.8–50% buffer B over a duration of 90 min with a flow rate of 300 nl per min. The Q-Exactive mass spectrometer (Thermo Fisher Scientific) was operated in data-dependent mode with survey scans acquired at a resolution of 70,000. Up to the top 10 most abundant from the survey scan were selected for fragmentation (nce=30). The maximum ion injection times for the survey scan and the MS/MS scans were 250 and 50ms respectively. The AGC target was set to 1E6 for survey scans and 1E5 for the MS/MS scans. MS/MS events were acquired at a resolution of 17,500. Dynamic exclusion was set at 20s.

### Protein identification and quantification

Relative protein quantitation was performed using Progenesis QI for proteomics (version 4.1, Nonlinear Dynamics) and the Mascot search engine (version 2.3.02, Matrix Science). Peptide spectra were searched against a Uniprot reference proteome (UP000002438, December 2016) and a contaminant database (cRAP, GPMDB, 2012) (combined 5733 sequences: 1,909,703 residues). The Mascot search parameters were as follows; precursor mass tolerance was set to 10 ppm and fragment mass tolerance was set as 0.01Da. Two missed tryptic cleavages were permitted. Carbamidomethylation (cysteine) was set as a fixed modification and oxidation (methionine) set as variable modification. Mascot search results were further validated using the machine learning algorithm Percolator embedded within Mascot. The Mascot decoy database function was utilised and the false discovery rate was set as <1%. Statistically significant differences in protein abundances between groups was determined by ANOVA analysis using Progenesis QI. Proteins with a q value ≤ 0.05 and a log2 fold change ≥ than (+/-) 1 were judged to be significant . Only proteins with ≥ unique peptides were used in any comparisons PCA plots were constructed using ClustVis (49).

### Kegg analysis

The proteins that satisfied the specified criteria were examined using the KEGG pathway mapper, where they were categorised into functional pathways. Subsequently, these pathways were visually examined using KEGG pathway mapper and then investigated using STRING (https://string-db.org/).

### Statistical analyses

The statistical analyses were conducted using Sigma Plot 14 software, unless specified otherwise. Growth was analysed using Growthcurver package in R (50) and biofilm assays were evaluated using Kruskal-Wallis analysis of variance and Holm-Sidak post hoc test for nonparametric data, unless otherwise stated. Visualisation was performed using R, ggplot2.

## Author statements

### Author contributions

H.E., funding, investigation, formal analysis, writing – original draft preparation; R.F., conceptualisation, writing – review and editing, supervision, project administration, analysis; NB., investigation, formal analysis; SA., proteomic analysis; AC., investigation, formal analysis; CB., investigation, formal analysis; J.F., conceptualisation, writing – review and editing, supervision, project administration, analysis.

## Abbreviations

AUM: Artificial Urine Media
CV: Crystal violet
CAUTI: Catheter associated infections
CF: Cystic Fibrosis
Escherichia coli: E. coli
Hcn: Hydrogen cyanide
Lysogeny Broth: LB
P. aeruginosa: Pseudomonas aeruginosa
PQS: Pseudomonas Quinolone Signal
Pch: Pyochelin
Pvd: Pyoverdine
UTI: Urinary tract infection.

