## Supplementary Figures S1-S5 for "Sex Hormones Alter *Pseudomonas aeruginosa* Iron Acquisition and Virulence Factors"

**Supplementary Information**

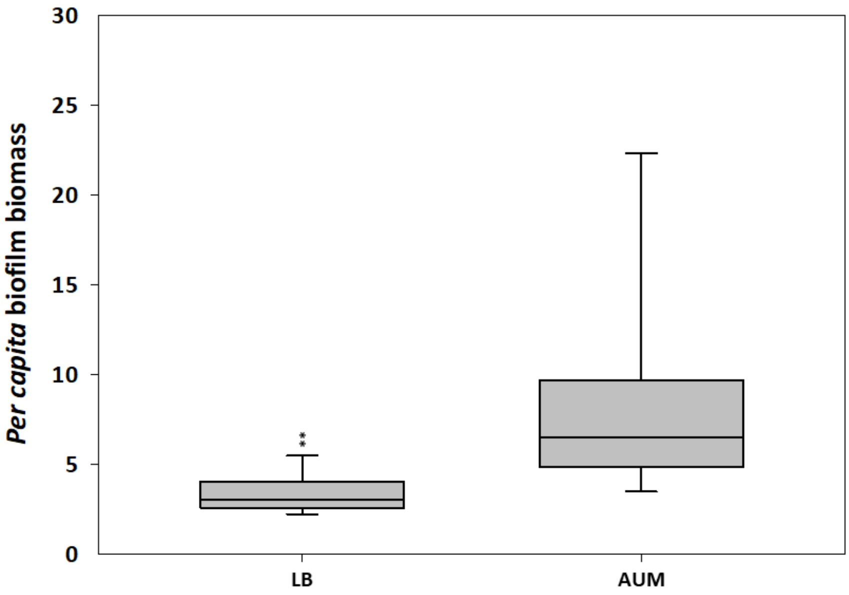


**Figure S1** Per capita biofilm biomass for 15 clinical *P. aeruginosa* clinical isolates in LB broth (LB) and artificial urine media (AUM) relative to $OD_{600}$ after 24 h. Significant differences are shown using **P<0.001.

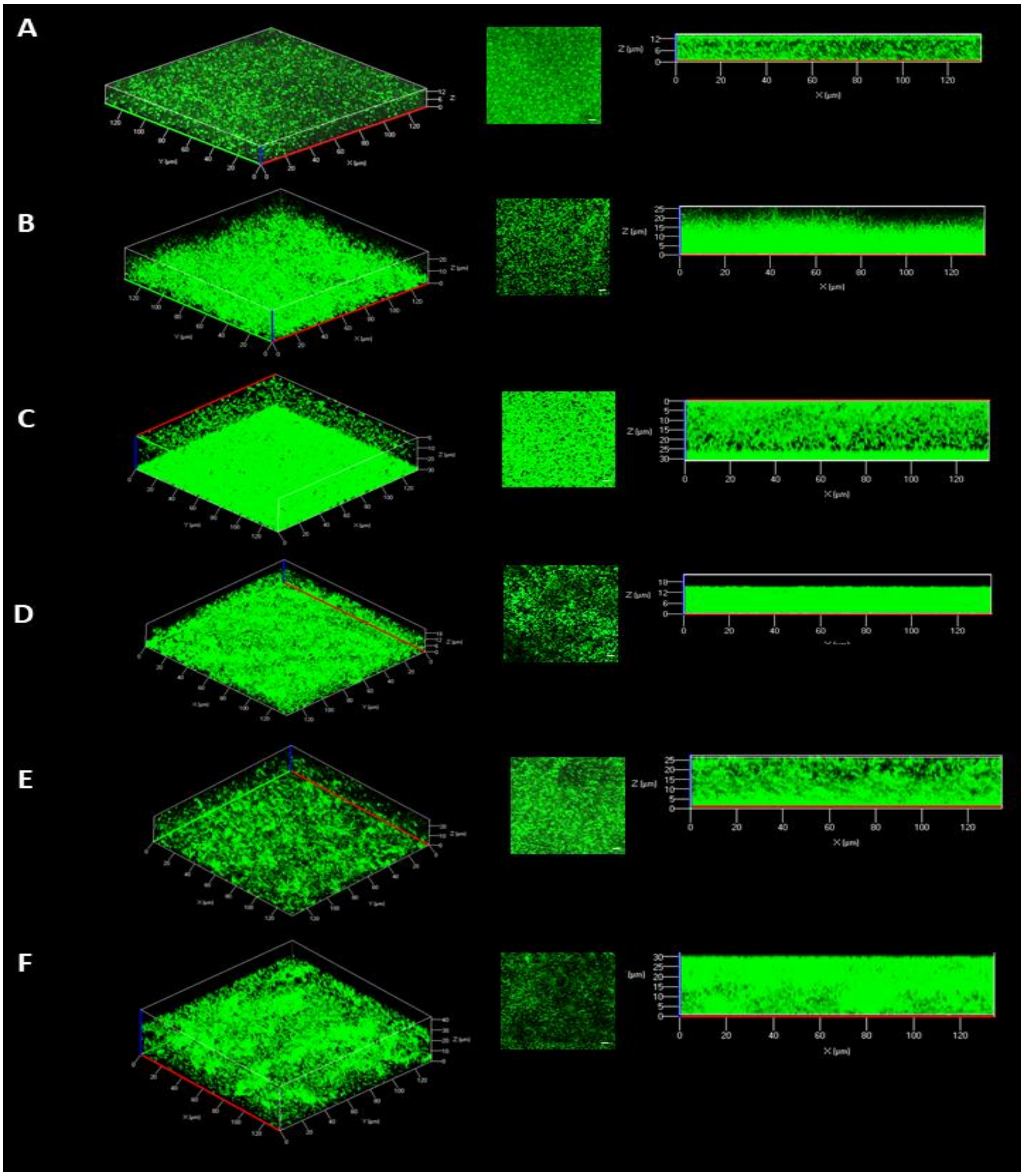


**Figure S2** Biofilm microscopy of *P. aeruginosa* grown in LB and AUM for 24h. Each panel contains an image of a 3D biofilm, an image of *P. aeruginosa* attached to the bottom of the chamber slide at the base of the biofilm and a cross section to display the profile of the biofilm. A) PAO1 grown in LB, B) PAO1 grown in AUM C) 133117 grown in LB, D) 133117 grown in AUM, E) 133043 in LB and F) 133043 in AUM. The white bar denotes 10 µm.

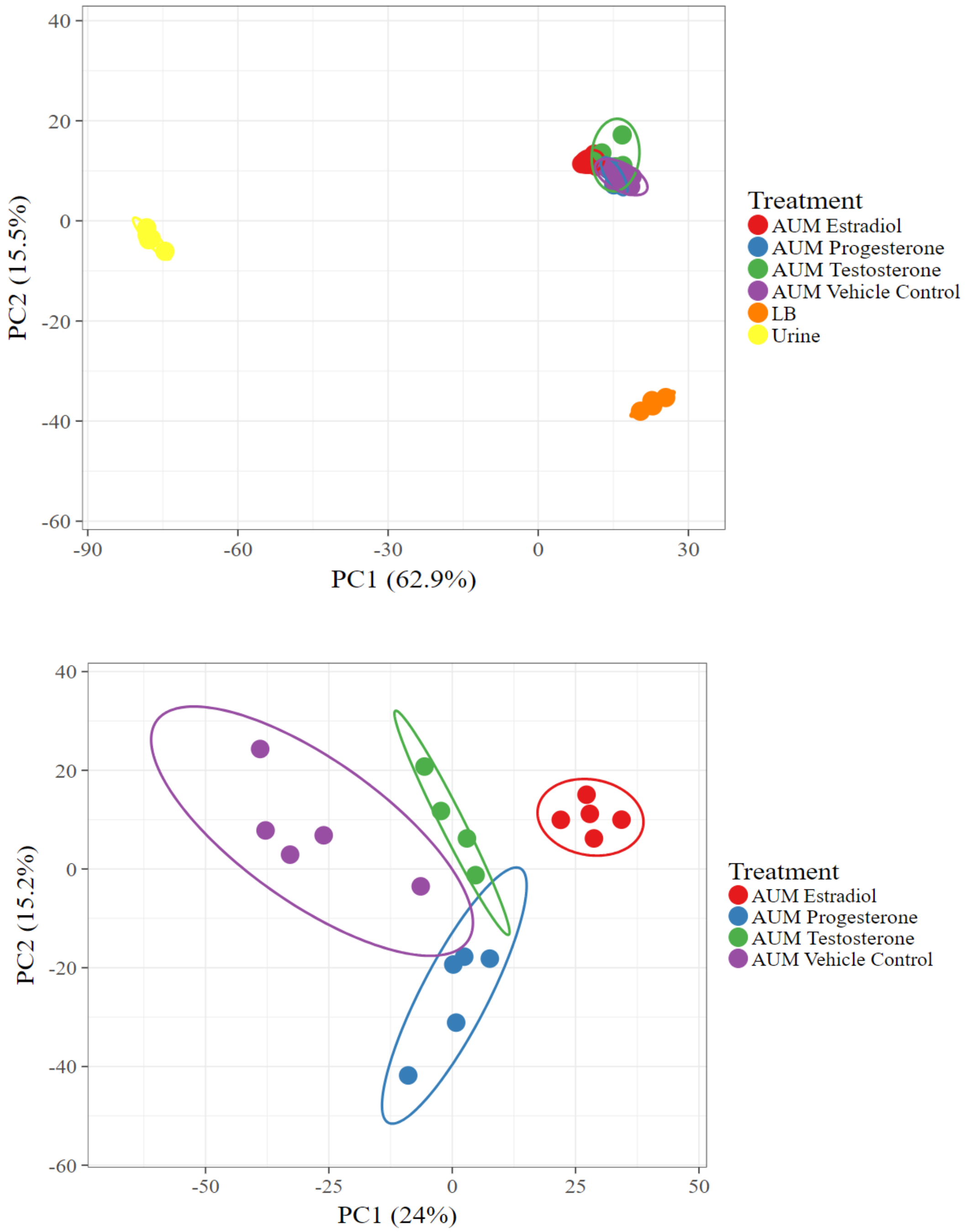


**Figure S3.** Principal component analysis of *P. aeruginosa* proteomes grown under different conditions. A) All *P. aeruginosa* proteome data showing growth in laboratory media (LB), artificial urine medium (AUM) and urine. Original values are ln(x + 1)-transformed. Unit variance scaling is applied to rows; SVD with imputation is used to calculate principal components. X and Y axis show principal component 1 and principal component 2 that explain 62.9% and 15.5% of the total variance, respectively. Prediction ellipses are such that with probability 0.95, a new observation from the same group will fall inside the ellipse. N = 29 data points. B). *P. aeruginosa* grown in the presence of sex hormones compared to an AUM vehicle control. Original values are ln(x + 1)-transformed. Unit variance scaling is applied to rows; SVD with imputation is used to calculate principal components. X and Y axis show principal component 1 and principal component 2 that explain 24% and 15.2% of the total variance, respectively. Prediction ellipses are such that with probability 0.95, a new observation from the same group will fall inside the ellipse. N = 19 data points.

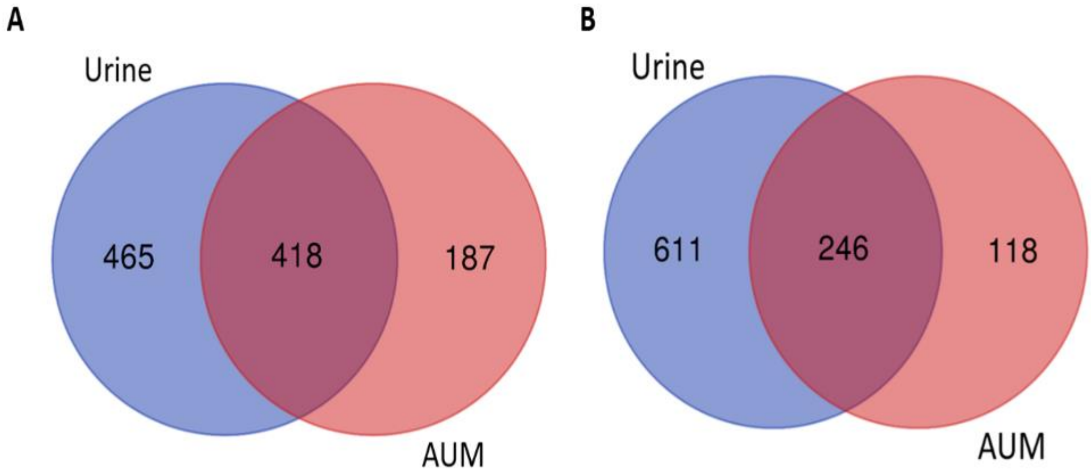


**Figure S4.** Altered *P. aeruginosa* protein abundance in urine and artificial urine medium (AUM). A) Increased abundance of proteins in hormones compared to LB. B) decreased abundance of proteins in hormones compared to LB.

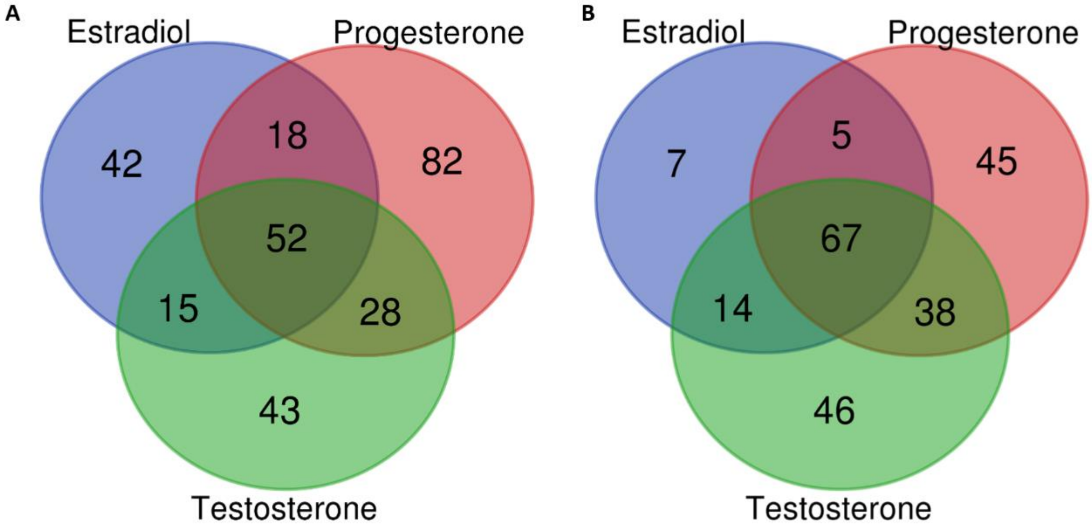

**Figure S5.** Altered *P. aeruginosa* protein abundance in the presence of sex hormones. A) Increased abundance of proteins in hormones compared to the vehicle control AUM-V. B) decreased abundance of proteins in hormones compared to the vehicle control AUM-V.
